# Feature representation under crowding in V1 and V4 neuronal populations

**DOI:** 10.1101/2021.10.21.465350

**Authors:** Christopher A Henry, Adam Kohn

## Abstract

Visual perception depends strongly on spatial context. A profound example is visual crowding, whereby the presence of nearby stimuli impairs discriminability of object features. Despite extensive work on both perceptual crowding and the spatial integrative properties of visual cortical neurons, the link between these two aspects of visual processing remains unclear. To understand better the neural basis of crowding, we recorded simultaneously from neuronal populations in V1 and V4 of fixating macaque monkeys. We assessed the information about the orientation of a visual target available from the measured responses, both for targets presented in isolation and amid distractors. Both single neuron and population responses had less information about target orientation when distractors were present. Information loss was moderate in V1 and more substantial in V4. Information loss could be traced to systematic divisive and additive changes in neuronal tuning. Tuning changes were more severe in V4; in addition, tuning exhibited greater context-dependent distortions in V4, further restricting the ability of a fixed sensory readout strategy to extract accurate feature information across changing environments. Our results provide a direct test of crowding effects at different stages of the visual hierarchy, reveal how these effects alter the spiking activity of cortical populations by which sensory stimuli are encoded, and connect these changes to established mechanisms of neuronal spatial integration.

---

Crowding is a perceptual phenomenon in which a clearly visible stimulus (a target) becomes difficult to discern when other stimuli (distractors) are presented in spatial proximity (Levi, 2008; Pelli and Tillman, 2008; Whitney and Levi, 2011; Manassi and Whitney 2018). Crowding has been studied extensively because it often limits visual function and because understanding its basis may provide insight into core aspects of visual processing.

Impairment of discriminability under crowding involves features from nearby stimuli being ‘combined’ with those of a target. The challenge for mechanistic descriptions of crowding is to identify the form this ‘combination’ takes at distinct stages of visual processing. There are numerous proposals of how crowding might arise, each invoking different computations and proposing different loci in the visual system. These proposals, some of which have conceptual overlap, include: lateral interactions between neurons within a cortical area that represent targets and distractors (Pelli, 2008; Levi and Carney, 2009); spatial pooling by neurons whose receptive fields encompass both the target and distractors (Levi, 2008); an encoding strategy in which visual stimuli are represented using a set of summary statistics (Balas et al., 2009; Freeman and Simoncelli, 2011) or simply averaged together (e.g. Parkes et al., 2001; Greenwood et al., 2009); or an erroneous readout of sensory information, involving either compulsory averaging or the substitution of signals encoding distractors for those encoding a target (e.g. Ester et al., 2015), perhaps arising from fluctuations in attention (He et al., 1996). These various bases of crowding have been proposed to be instantiated in primary visual cortex (V1; Levi and Carney, 2009; Pelli, 2008), V2 (Freeman and Simoncelli, 2011; He et al., 2019), V4 (Levi, 2008), or a combination of these (Freeman et al., 2011b).

To date, how crowding affects neural representations has been studied almost exclusively using coarse or indirect measures of brain activity, such as EEG or functional MRI (Bi et al., 2009; Anderson et al., 2012; Millin et al., 2014; Chen et al., 2014; Kwon et al, 2014; Chicherov et al., 2014). These approaches provide important information, but they lack the resolution to elucidate how crowding affects the representation of visual information by neuronal spiking activity, the activity that encodes and transmits visual information in the brain. As a result, we have a limited understanding of how crowding arises.

We recently assessed the degree to which crowded visual displays limit neuronal population information about the orientation of a target stimulus, in the primary visual cortex (V1) of anesthetized macaque monkeys (Henry and Kohn, 2020). We found that distractors reduced information through a combination of divisive, suppressive modulation of responsivity in some neurons and additive facilitation in others. The information loss in V1 could account for much, but not all, of the perceptual crowding evident in humans and monkeys performing a discrimination task using the same stimuli. Thus, we proposed that spatial contextual modulation in V1 is a major contributor to crowding.

How crowded displays influence stimulus encoding in midlevel visual cortex is unknown. The more extensive spatial pooling performed by neurons in higher visual cortex (Van Essen et al., 1984; Gattass et al., 1988) suggests additional mechanisms by which sensory information may be lost, though precisely how those neurons pool visual inputs across space is not well understood (Bushnell et al., 2011; Motter, 2018). Here we assess how information about target orientation is affected by the presence of distractors in neuronal populations in area V4 of fixating macaque monkeys. We find that V4 populations show greater information loss than in simultaneously recorded V1 populations, under identical stimulus conditions. Further, this loss is evident for a greater range of spatial offsets between targets and distractors. Information loss in V4 involves some of the same mechanisms previously seen in V1, but V4 neurons also show a complex dependence on the precise set of stimuli presented in or near their spatial receptive fields.

## Results

We carried out simultaneous extracellular recordings from V1 and V4 neuronal populations via chronic microelectrode arrays implanted in two awake macaque monkeys (Figure 1A). Animals viewed stimuli passively, while maintaining fixation within a 1.3 degree diameter window about a small central point. We first obtained maps of the spatial receptive fields of the sampled neuronal populations in dedicated recording sessions. As expected, individual V1 receptive fields were small, and the receptive fields of V4 neurons were many-fold larger (Figure 1B, top). Aggregate receptive field maps showed substantial overlap in the visual field representation of recorded V1 and V4 populations in each animal (Figure 1B, bottom).

**Figure 1.**
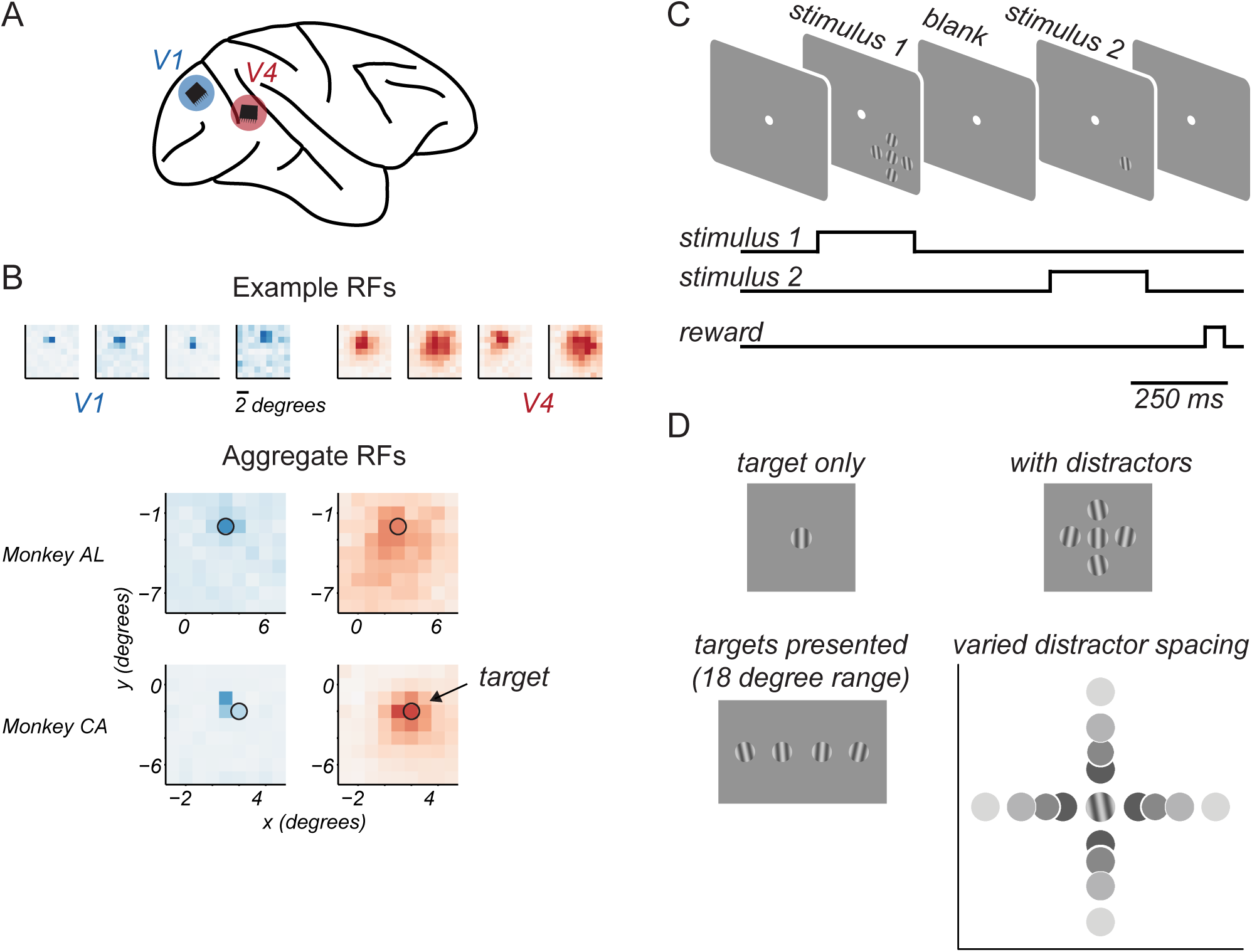
Experimental approach. A. Microelectrode arrays were chronically implanted into V1 and V4 of one hemisphere of each animal. B. Top. Spatial receptive field maps of 4 example neurons from V1 (blue) and V4 (red). Darker colors indicate higher firing rates. Bottom. Aggregate receptive field maps of recorded neuronal populations, by cortical area (columns) and animal (rows). Black circle indicates target stimulus location and approximate size. C. Task structure.Animals maintained fixation of a central spot while target stimuli were presented in two 0.25s epochs on each trial, separated by a 0.35s blank period. D. Target stimuli were drifting gratings chosen from an 18 degree range around a reference orientation. When shown, distractors were presented at ±10 deg relative to this reference orientation. Target-distractor spacing was varied.

To assess feature encoding, we used a small target grating (1 degree diameter), positioned to drive both cortical populations. The target grating was located at 2.8 degrees eccentricity in the lower visual field of one animal, and at 3.6 degrees in the other (indicated by the black circle, Figure 1B). On each trial, stimuli were shown in two epochs separated by 0.35 s (Figure 1C). Target stimuli were chosen from an 18-20 degree range straddling a reference orientation (changed in each session), in increments of 4-6 degrees (Figure 1D). On each presentation, the target stimulus could be shown either in isolation or surrounded by 4 distractor stimuli, whose orientation was ±10 degrees of the reference orientation. The spacing of distractors around the target was varied because target-distractor distance is known to influence perceptual crowding (Bouma, 1970). Identical stimuli, save for a rotation in reference orientation by 90 degrees, were shown on half the presentations, to minimize the effects of adaptation (Kohn, 2007). Thus, each recording session probed cortical responses to two narrow ranges of target orientation, yielding distinct responses from the recorded neuronal populations that were analyzed independently.

We recorded in 11 sessions, yielding 22 neuronal populations in each area. The size of V1 populations was on average 19 ± 1.9 neurons, and the size of V4 populations was 31 ± 3.4 neurons.

### Distractors limit target discriminability

We first assessed how the presentation of distractors affected the discriminability of target orientation afforded by responses of individual V1 (n = 396) and V4 neurons (n = 691). To quantify discriminability, we used receiver operating characteristic (ROC) analysis to distinguish between the spike counts evoked by two target gratings differing 18 degrees in orientation (Figure 2A, abscissa). Chance performance is indicated by a value of 0.5 in the area under the ROC curve; values of 0 or 1 indicate perfect discriminability between stimuli (for neurons with opposing tuning slopes over the sampled target range). Note that performance in many neurons was near chance, because the narrow range of target orientations did not include those that modulated the neuron’s responses significantly (i.e. near the flank of the tuning function).

**Figure 2.**
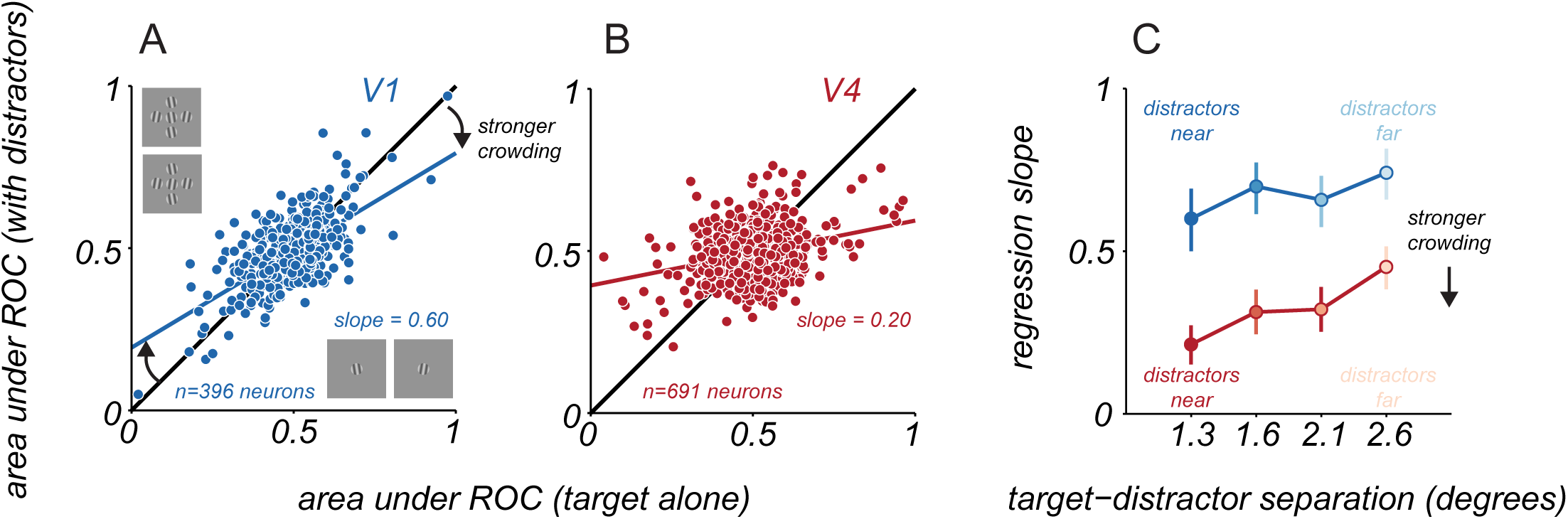
Single neuron discriminability of target orientation. A. Target discriminability of V1 neurons for targets alone (abscissa) and targets with near distractors (ordinate), using responses to the two most differing target orientations. Solid blue line indicates linear regression fit to the data. B. Same as A, but for V4 neurons. C. Change in discriminability as a function of target-distractor spacing for V1 (blue) and V4 (red), quantified using the slope of the regression line. Error bars indicate 95% confidence interval.

We compared single neuron discriminability for targets alone and targets with distractors. For V1 neurons, discriminability was better when targets were presented alone (Figure 2A), evident as a regression slope that was significantly less than 1 (slope = 0.60, 95% confidence interval: [0.50 0.69]). When distractors were moved further from the target, effects on discriminability were slightly more moderate (Figure 2C, farthest spacing, slope = 0.74, [0.66 0.82]). In V4, the loss of discriminability with nearby distractors was stronger (Figure 2B, slope = 0.20, [0.14 0.26]) and remained pronounced even when distractors were placed farther from the target (Figure 2C, farthest spacing, slope = 0.44, [0.37 0.50]). Thus, distractors led to a greater impairment in single neuron discriminability of targets in V4 than V1, over all spatial scales.

We next quantified how distractors affected neuronal population information about target orientation. We used linear discriminant analysis to determine how well two target stimuli could be distinguished using the measured neuronal population responses. We optimized decoders separately for each pairing of target stimuli, when targets were presented alone or with distractors. By allowing for separate decoding of these sets of responses, we provide an upper-bound estimate on the target information accessible to a linear readout strategy of the measured responses.

To provide good estimates of how distractors affect representational accuracy, we aggregated decoding performance across target grating pairings and sessions from each animal, for each cortical area and target-distractor spacing. An example is shown in Figure 3A,B for V1 and V4 of one animal, comparing performance for responses to targets alone and to targets with the nearest distractors. When decoder performance was above chance (abscissa values greater than 0.5), performance for responses to targets with distractors tended to be lower (ordinate). The running average of decoding performance for responses with distractors as a function of target only performance (blue and red lines, in Figures 3A and 3B respectively) were consistently below the diagonal, indicating performance was worse with distractors.

**Figure 3.**
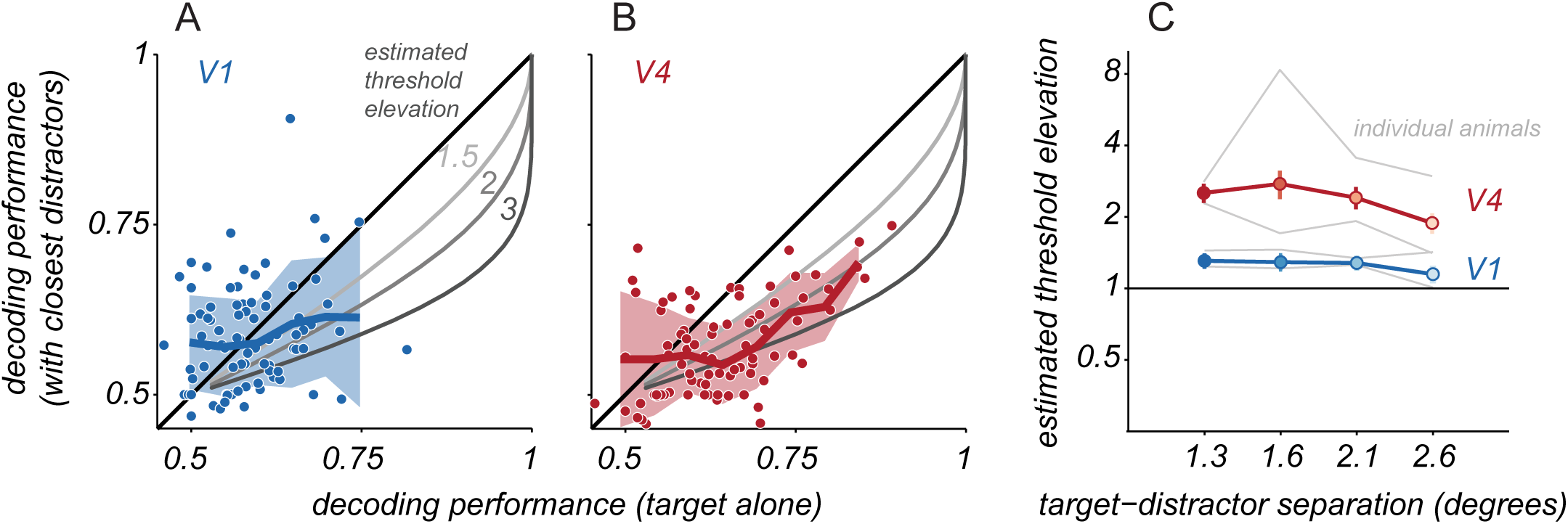
Neuronal population target discriminability. A. Cross-validated V1 population discrimination performance for targets alone (abscissa) and targets with distractors (ordinate). Each point represents a single pairing of two target stimuli. Data are combined across pairings and sessions, for one animal and the nearest target-distractor spacing (n=84 cases). Solid blue line and shaded region represent running average and standard deviation of crowded performance. Gray lines indicate expected performance for the indicated threshold elevations. B. Same as A. but for V4. C. Estimated threshold elevation for populations as a function of target-distractor separation. Points represent the mean across both animals (V1, blue; V4, red); error bars indicate s.e.m., estimated via bootstrapped resampling of the data. Gray lines represent the mean values for each animal and cortical area.

To estimate how the change in decoder performance might translate to the strength of perceptual crowding, we fit a single parameter model to the decoder performance values aggregated across sessions. The model consisted simply of a psychometric function for target discrimination whose slope could change with the appearance of distractors. The single parameter provided an estimate of the discrimination threshold elevation that would be consistent with the change in population performance (see Figure 3 – figure supplement 1). Sample predicted performance trends for different changes in threshold are shown in Figures 3A,B (gray curves).

In V1, distractors led to a loss of population information about target orientation whose magnitude was consistent with a moderate elevation in discrimination threshold (Figure 3C, blue). Distractors at the closest spacing yielded the largest threshold elevations, 1.31 ± 0.10, and distractors at the farthest spacing yielded the smallest elevation, 1.15 ± 0.08. The estimated threshold elevation was statistically significant for all but the farthest target-distractor separation (permutation test, p<0.05). In V4 populations, the estimated threshold elevations were greater than those in V1 (Figure 3C, red): 2.52 ± 0.23 for the closest distractors and 1.89 ± 0.19 for farthest distractors (significant for all distractor separations, permutation test, p<0.001).

We were concerned that our estimates of threshold elevation might be poorly constrained by the limited decoding performance values, which reflected the small size of the recorded populations. We thus repeated our analysis using responses of larger ‘pseudo-populations’, constructed by combining neurons across sessions. Pseudo-populations provided a broader range of performance values, and yielded similar estimates of how distractors affected thresholds (Figure 3 – figure supplement 2).

In summary, we found that distractors reduced population information about target orientation in both V1 and V4. In V1, there was a moderate threshold elevation for the closest distractors and little effect for distractors placed farther away. In V4, there was a much greater threshold elevation, evident across a wider range of distractor spacings. We next turn to describing which changes in neuronal activity contribute to the loss of encoded feature information.

### Distractors modulate neuronal responsivity and variability

The stimulus information encoded by neuronal populations (as extracted by linear decoders) is a function of the tuning and response variability of the individual neurons and of how their variability is shared (Averbeck et al., 2002; Kohn et al., 2016). All of these response statistics can be altered by changes in spatial context, such as the presentation of distractors (Henry et al., 2013; Snyder et al., 2014; Trott and Born, 2015; Henry and Kohn, 2020). To understand why V1 and V4 neuronal populations provide less information about target orientation in crowded displays, we assessed how distractors affected pairwise correlations of variability, neuronal variability, and neuronal responsivity, in turn.

We first determined whether the presence of distractors altered the strength of shared variability among neurons. Distractors significantly reduced pairwise ‘noise’ correlations (r_SC_) compared to those evident in responses to targets alone. In V1 (Figure 4A), pairwise correlations decreased from 0.07 ± 0.003 for targets alone to 0.05 ± 0.002 with the nearest distractors (paired t-test, p<0.001). In V4 (Figure 4B), the corresponding decrease in correlations was from 0.11 ± 0.001 to 0.06 ± 0.001 (paired t-test, p<0.001). To determine whether altered correlations contributed to information loss, we fit decoders to population responses after trial shuffling responses to reduce correlations. In these shuffled data, distractors led to threshold elevations similar to those seen in the original data (Figure 4C), indicating changes in correlations, though substantial, played a minimal role in the information loss we observed.

**Figure 4.**
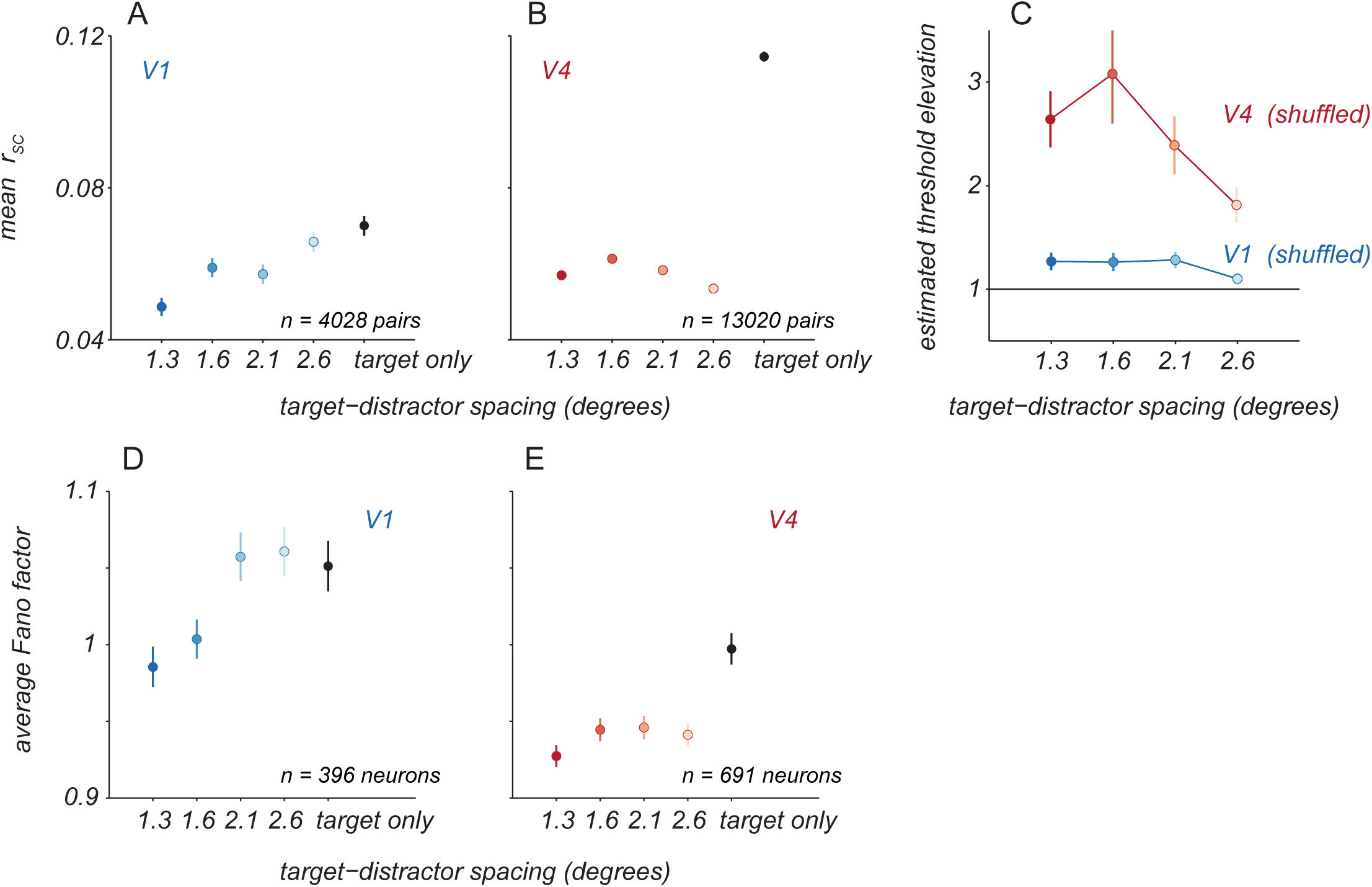
Effect of distractors on neuronal variability. A. Average pairwise ‘noise’ correlations (rSC) for V1 neuronal pairs for targets alone (black) and with distractors of varied spacing (blue). Points and error bars represent mean ± s.e.m. B. Average correlations for V4 neuronal pairs, same conventions as in A. C. Crowding effect size for V1 and V4 populations when decoding was performed on trial-shuffled population activity. Points and error bars represent mean ± s.e.m. D. Individual neuron variability was quantified as the average Fano factor for V1 neurons for targets alone (black) and with distractors at various spacings (blue). Points and error bars represent the geometric mean ± s.e.m. over all neurons. B. The average Fano factor for V4 neurons, same conventions as in A.

To quantify how response variability was affected by distractors, we measured the response variance normalized by its mean (Fano factor). The Fano factor for each neuron was calculated as the geometric mean of the Fano factors measured for each target orientation separately. In V1, distractors at the closest spacing led to a slight reduction in variability (Figure 4D, from 1.05 ± 0.02 for targets alone to 0.98 ± 0.01, paired t-test, p<0.001). In V4, distractors at all spacings led to a significant reduction in variability (Figure 4E, 1.00 ± 0.01 for targets alone to 0.93 ± 0.01 with closest distractors, paired t-test, p<0.001 for each spacing). All else being equal, the reduction in variability with distractors should result in better discriminability not worse. Thus, changes in variability must not be responsible for information loss we observe.

We next asked how distractors influenced neuronal responsivity. We calculated a modulation index defined as the ratio of the response to targets with distractors relative to that for targets alone. For each neuron, we summarized this index across targets as the geometric mean of the indices computed for each target orientation separately. In V1, neurons exhibited diversity of modulation, with both suppression and facilitation evident. For distractors at the closest spacing, modulation was slightly suppressive on average (Figure 5A, dark blue points, 0.98 ± 0.44, median ± s.d.). For distractors farther from the target, facilitation became less prevalent and modulation remained slightly suppressive on average (Figure 5A, light blue points; 0.96 ± 0.07). The responses of V4 neurons could also be facilitated or suppressed by the presentation of distractors. For the closest distractors, most V4 neurons showed facilitation (Figure 5B, dark red points, 1.11 ± 0.38, t-test, p<0.001) and the average modulation index was larger than in V1 (one sided t-test, p<0.01). As distractor spacing increased, a roughly equal proportion of V4 neurons exhibited facilitation and suppression (Figure 5B, light red points, 0.97 ± 0.48).

**Figure 5.**
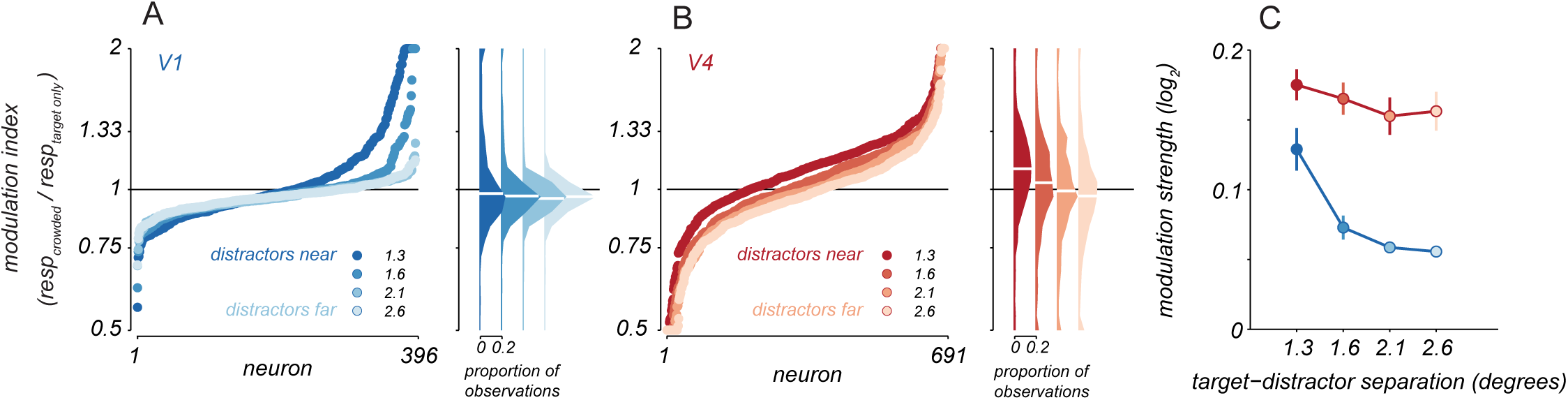
Response modulation with distractors. A. Modulation indices were calculated as the average response to targets with distractors over the response to targets alone. Points represent sorted modulation indices for all V1 neurons, with target-distractor spacing indicated by blue shading. Distributions of modulation indices are shown at the right, with median values for each distribution indicated by white lines. B. Modulation indices for V4 neurons in red, same conventions as in A. C. Average modulation strength as a function of target-distractor spacing for V1 (blue) and V4 (red). Points represent mean ± s.e.m (V1: n=396, V4: n=691 neurons).

To quantify the strength of modulation by distractors, regardless of whether it facilitated or suppressed responsivity, we averaged the absolute value of each neuron’s modulation index (log-transformed). In V4, the average modulation was stronger than in V1 at all separations (Figure 5C, one sided t-test, p=0.01 for closest distractors and p<0.001 for all other spacings). Distractors at the farthest separation yielded weak modulation in V1 (one sided t-test, p<0.001) but modulation strength in V4 was largely maintained (one sided t-test comparing to nearest distractors, p=0.44). Thus, while the nature of modulation in V4 changed with the spatial scale of visual stimulation, the strength of modulation was similar across the range of target-distractors spacings explored.

To understand how response modulation by distractors influences the discriminability of target orientation, we must consider the functional form of changes in orientation tuning induced by distractors. We therefore fit a model in which each neuron’s target tuning amid distractors was described by a multiplicative scaling and additive translation of the target alone tuning (Figure 6A, shaded regions). The additive term was normalized, so that it reflected the proportion of the mean firing rate for targets alone that was needed to produce the appropriate translation in tuning. In this analysis, we only considered neurons that had reliable tuning over the narrow range of sampled orientations (see Methods).

**Figure 6.**
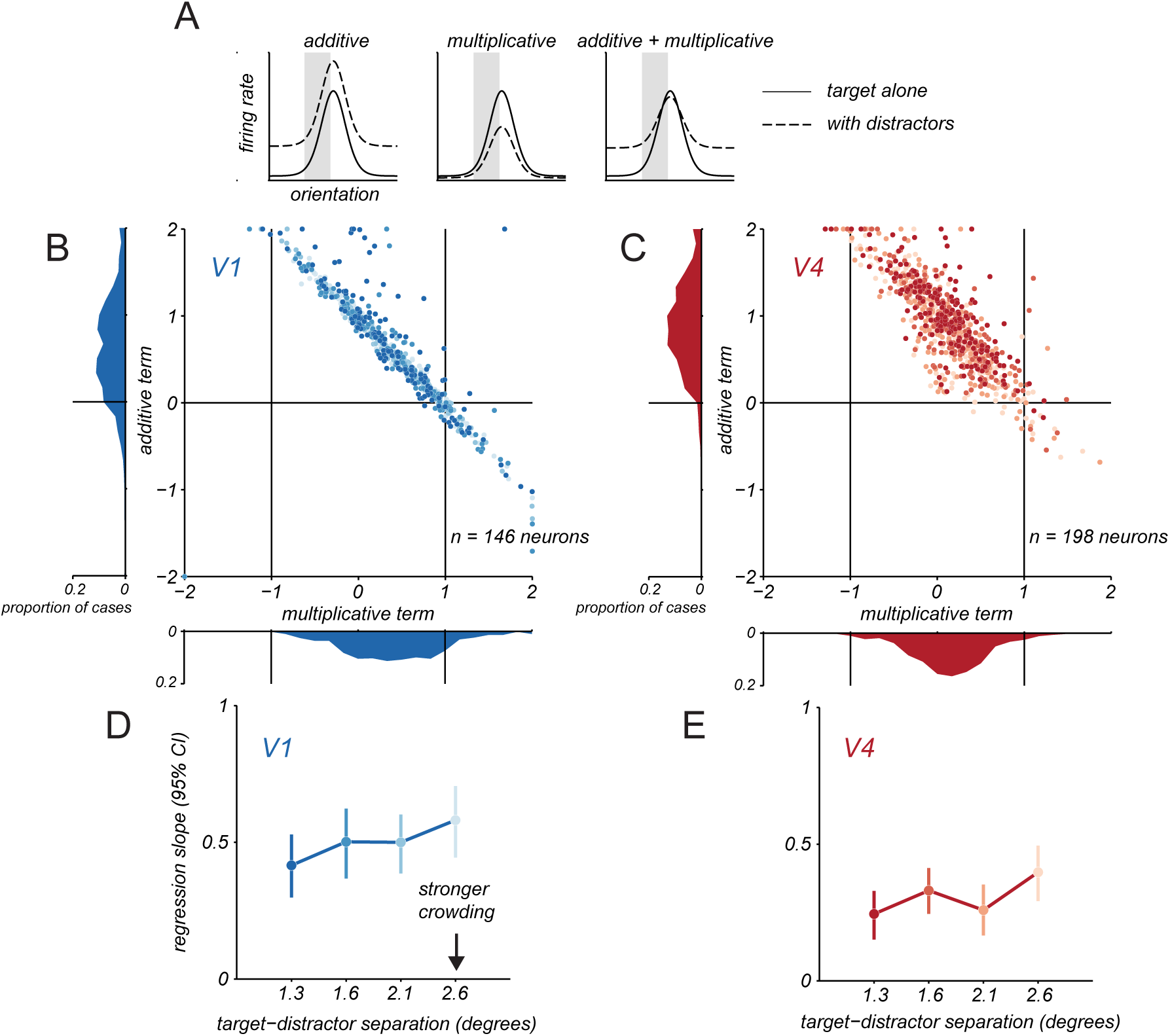
Additive and multiplicative changes in target tuning with distractors. A. Schematic illustration of how target tuning amid distractors (dashed line) might involve additive translation, multiplicative scaling, or combined changes in target only tuning (solid line). B. Multiplicative and additive terms fit to V1 neurons under crowding. Each point represents one neuron and distractor spacing condition, darker shades indicate closer distractor spacing. Marginal distributions of parameters are shown along the axes. C. Same as in B, but for V4 neurons. D. Change in V1 discriminability as a function of target-distractor spacing by applying the additive and multiplicative changes in target only tuning from B. Change in discriminability was quantified using the slope of the regression line relating discriminability for target only and target with distractors conditions. Error bars indicate 95% confidence interval. E. Same as in D, but for V4.

Figure 6 shows the fitted parameters for V1 (Figure 6B) and V4 (Figure 6C) neurons, with each point representing a neuron and distractor spacing condition, and darker points representing closer distractors. By definition, there was strong covariation between the additive and multiplicative terms. For tuning that became flat with distractors, the multiplicative term was 0 and the additive term was positive to reproduce the mean firing rate. For tuning that was only partially flattened, the multiplicative term was more positive (but less than 1), so a smaller additive term was required to sum to reproduce the tuning curve. When tuning slope reversed in the presence of distractors, the multiplicative term was negative; as a result, a large additive term was required to offset the negative responses and reproduce the firing rate. Note that these reversals in tuning slope occurred rarely in V1 but more often in V4; such striking changes in tuning are considered in the following section.

For V1 neurons (Figure 6B), the amplitude of the multiplicative term was significantly less than 1 on average (abscissa, 0.59 ± 0.03), indicating a compression of the tuning curve, and the additive term was positive (0.39 ± 0.16), indicating an upward translation of the tuning. For V4 neurons (Figure 6C), the amplitude of the multiplicative term was also less than 1 (0.36 ± 0.01) and the additive term was positive (0.90 ± 0.02). The multiplicative term for V4 neurons was significantly smaller than for V1 neurons (t-test, p<0.001) and the additive term was significantly larger for V4 than V1 (t-test, p>0.001). In summary, the presence of distractors compressed the dynamic range of neuronal tuning in both areas, via a multiplicative scaling, and maintained or enhanced responsivity, via an additive offset.

Both the compression of dynamic range and the additive shift in responsivity would be expected to reduce discriminability. The former would make responses to different stimuli more similar; the latter would result in greater response variability, since variability in cortical responses scales with the mean. To test whether these two effects capture the information loss we observe in V1 and V4, we simulated crowded responses by scaling and translating the target alone tuning with the fitted multiplicative and additive terms. We then calculated the discriminability for targets alone and for targets with distractors, using the area under the ROC curve (as in Figure 2). This simulation yielded some loss of discriminability in V1 (Figure 6D) and a more substantial loss in V4 (Figure 6E). In both areas, the loss of discriminability depended weakly on target-distractor spacing, as in the data. In additional analysis, we considered an alternate simulation allowing only for divisive modulation in suppressed neurons and additive modulation in facilitated neurons (Figure 6 – figure supplement 1), as in Henry and Kohn (2020). Both these forms of modulation reduced discriminability, but understated the strength of observed effects, particularly in V4.

Together, these results demonstrate that divisive scaling and additive translation of target tuning in the presence of distractors limit the encoding of target orientation, and that additional impairment in V4 results from greater modulation by these factors compared to V1.

### Distractors alter neuronal tuning for target orientation

Both additive facilitation and divisive suppression of neuronal responsivity by distractors are consistent with well-established mechanisms of spatial integration by cortical neurons (Movshon, 1978; Carandini et al., 2005; Carandini and Heeger, 2011). Yet, previous work has also suggested more complex forms of spatial integration, particularly in V4. For instance, the tuning of V4 neurons for a small stimulus in the receptive field can change dramatically, and often seemingly idiosyncratically, when paired with other stimuli (e.g., Bushnell et al., 2011; Motter, 2018). Consistent with this description, we often observed that V4 tuning for target orientation could be altered substantially when distractors were present (Figure 7B). V1 tuning for targets, in contrast, was usually (Figure 7A, left) but not always (right) scaled or translated by distractors.

**Figure 7.**
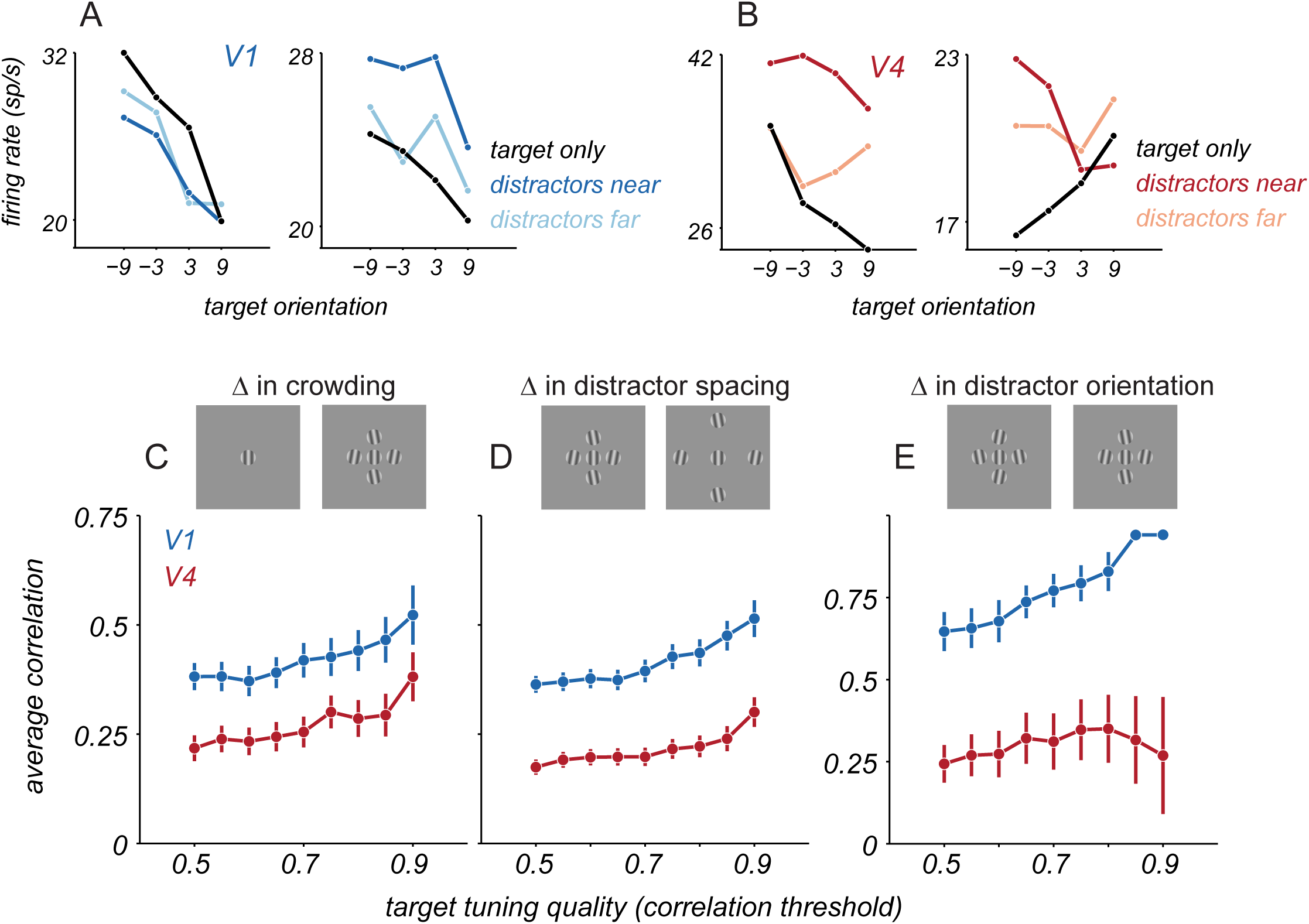
Change in target tuning with distractors. A. Target orientation tuning for 2 example V1 neurons for targets alone (black) and for targets with near (dark blue) or far (light blue) distractors. B. Target orientation tuning for 2 example V4 neurons for targets alone (black) and with distractors (red). C. Average correlation between tuning for targets alone and targets with near distractors for V1 (blue) and V4 (red), as a function of target tuning quality. Points and error bars represent mean ± s.e.m. D. Average correlation between target tuning with near and far distractors. E. Average correlation between target tuning for two near distractor configurations, counterbalanced for distractor orientation.

To quantify the similarity of tuning for targets alone and targets with distractors, we calculated their Pearson correlation. If target tuning is simply scaled or translated by distractors, correlation values should be high. If distractors distort tuning, correlations should be lower. To provide a fair comparison of V1 and V4 tuning, we selected subsets of neurons in each area that exceeded a criterion that captured the reliability of tuning for targets alone, defined by calculating the correlation between the tuning from even versus odd trials. In V1, tuning for targets alone was well correlated with the tuning for targets with nearby distractors (Figure 7C, blue points). For instance, for cells exceeding a criterion of 0.5, the correlation between tuning with and without distractors was 0.38 ± 0.03. Tuning correlation was significantly lower in V4 neurons (Figure 7C, red points), with the corresponding value being 0.22 ± 0.03 (p<0.001 for comparison with V1, one-sided t-test).

We next assessed whether the change in tuning depended on the particular set of distractor gratings. We computed the correlation of target tuning for displays with different target-distractor spacings (near and far). In this comparison too, correlations between target tuning were lower in V4 than V1 (Figure 7D, for cutoff of 0.5, correlations were 0.17 ± 0.02 and 0.36 ± 0.02, respectively, p<0.001, one-sided t-test). Even subtle changes in the composition of distractors could strongly alter tuning for the target. Specifically, we compared target tuning in the presence of two distractor configurations, for which the position of each distractor was fixed but its orientation differed by 20 degrees between configurations (with the same average orientation within each configuration). V4 tuning differed substantially between these two conditions, whereas V1 tuning was nearly unaffected (Figure 7E, for cutoff of 0.5, V4 correlation values were 0.24 ± 0.06 and V1 correlation values were 0.65 ± 0.06, p<0.001 for difference between areas, onesided t-test). Together, these results suggest that distractors introduce configuration-dependent changes in neuronal tuning for target orientation, particularly in V4.

The change in V4 tuning with distractors need not result in a loss of population information about target orientation, because changes in tuning could be compensated for by adjusting the readout strategy. Indeed, the population decoding we have considered thus far had this flexibility, as it involved fitting separate decoders to the activity evoked by each display. However, circuits reading out V1 and V4 responses are unlikely to flexibly alter their strategy, to optimally decode responses for each stimulus condition. For a fixed decoding strategy, changes in tuning with distractors, as described above, would be expected to limit discriminability further. To assess this possibility, we projected population responses onto two fixed decoding axes: one that was optimized for responses to targets alone and one optimized for targets with nearby distractors.

Figure 8A shows example population responses from a single V1 session projected onto these two decoding axes, with ellipses indicating response distributions for each of the 4 target orientations. In V1, projection of responses from either condition yielded similar separation of points on either decoding axis, indicating the information about target orientation can be decoded similarly well by a fixed decoder optimized either to responses to targets alone or to targets with distractors. In V4 (Figure 8B), projections were less aligned between the two decoding strategies, and the separation of points was reduced along the decoding axis of the opposing condition. Thus, in V4, projections depend heavily upon the decoding axis used, and decoding performance would be expected to generalize poorly between configurations.

**Figure 8.**
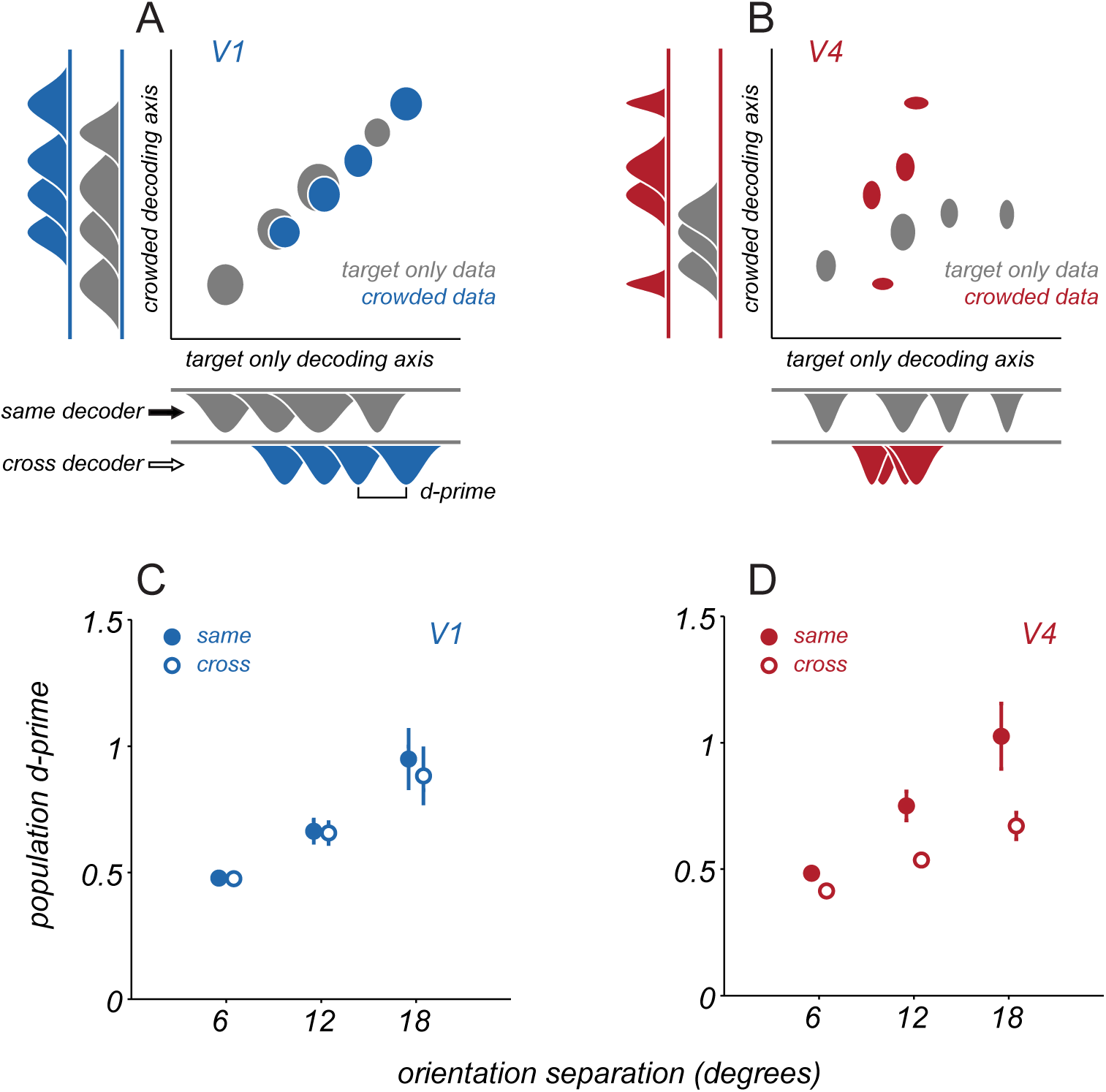
Generalization of discriminability across decoding axes. A. An example V1 population response projected onto two distinct decoding axes, one optimized for target only responses (abscissa) and one optimized for responses to targets with distractors (ordinate). Gray ellipses represent mean ± s.e.m. over repeated trials of a single target stimulus, with the 4 ellipses representing the distinct target orientations shown. Blue ellipses represent the same, but for responses to targets with distractors. Marginal distributions are shown at the sides of the plot. B. An example V4 population response, same conventions as in A. C. Average V1 discriminability is measured as the d-prime between two distributions, when projected onto the same decoding axis as the data (same, filled circles) or the decoding axis from the opposing condition of crowding (cross, open circles). Points and error bars represent mean ± s.e.m. Points are offset slightly along abscissa for visualization. D. Average V4 discriminability, same conventions as in C.

To quantify how well a common decoding strategy worked across configurations, we calculated the dprime value between response distributions for each pair of target stimuli, when projected onto a decoding axis defined by a matching display condition (same projection) or the opposite configuration (cross projection). Discriminability between stimuli was high in V1 and V4 when using the matched decoder (‘same’, Figures 8C,D). When data were projected onto the decoding axis of the opposing condition (‘cross’), discriminability was largely preserved in V1 (dprime of 0.95 ± 0.13 for same projection and 0.88 ± 0.12 for cross projection, p=0.20, paired t-test). In contrast, discriminability dropped significantly for V4 (1.02 ± 0.13 vs. 0.67 ± 0.05, p=0.002, paired t-test). In additional analyses, we confirmed that the distinct decoding axes in V4 for different displays do not arise from relying on distinct subsets of neurons. (Figure 8 – figure supplement 1). Thus, changes in V4 representation across displays do not result from dramatic changes in which neurons were relevant for encoding target orientation, but rather from changes in the tuning of a common underlying pool of neurons between conditions.

Together, our results indicate that V4 populations exhibit a more configuration-dependent representation of target orientation than that in V1. If perception involves a fixed linear readout of V4 responses for different displays, discriminability is likely to be strongly compromised.

## DISCUSSION

We evaluated how neuronal population information about target orientation in areas V1 and V4 is affected by the presence of distractors. V1 (Levi and Carney, 2009; Pelli, 2008; Chen et al., 2014; Kwon et al., 2014; Millin et al., 2014) and V4 (Motter 2018; Levi, 2008) have each been suggested as a possible locus for the bottleneck limiting perception in crowded scenes. We found that distractors cause a moderate loss of information about target orientation in V1 and a more pronounced loss in V4. V4 effects extended over a larger range of target-distractor separations than V1 effects. Distractors affected the responsivity, tuning, and variability of neurons in both areas. Responses were diversely modulated, but modulation often took the form of a divisive suppression and additive facilitation of responsivity, whose combined influence could explain the reduction in discriminability with distractors. Our results provide a direct test of crowding effects at different stages of the visual hierarchy using neuronal population spiking activity and connect changes in representational quality to established mechanisms of neuronal spatial integration.

Both facilitation and suppression of responses to a target by distractors are consistent with prior descriptions of how V1 and V4 neurons integrate visual signals spatially. Response suppression in V1 is expected: distractors fell in the receptive field surround of most of the sampled neurons, and stimuli in the surround often elicit suppression that acts divisively (Cavanaugh et al., 2002; Carandini and Heeger, 2011). Facilitation in V1 would be expected in those neurons whose receptive fields straddle target and distractor stimuli (Movshon et al., 1978; Cavanaugh et al., 2002). In V4, neuronal receptive fields are a few-fold larger than in V1, at matched eccentricity (Gattass et al., 1988), and thus more likely to span multiple elements of crowded displays. Within the classical receptive field, V4 neurons have been shown to sum multiple visual inputs (Ghose and Maunsell, 2008), as well as to show various forms of normalization (Verhoef and Maunsell, 2016). In addition, V4 neurons possess a suppressive surround (Zanos et al. 2011; Ni et al. 2012; Verhoef and Maunsell, 2016) that would be expected to be engaged by the distractors in our displays, particularly when they are far from the target.

We have previously shown that discriminability in neurons with Poisson-like variability can be reduced either if distractors recruit divisive suppression or additive facilitation of tuning (Arandia-Romero et al., 2016; Henry and Kohn, 2020). We also reported that, in V1, suppressed neurons were better explained by a divisive modulation, and facilitated neurons better explained by an additive modulation. We replicated those findings for the V1 and V4 populations in the present study (Figure 6 – figure supplement 1A-B). However, as is clear from Figure 6, most neurons in V1 and V4 are better described as having both multiplicative and additive changes in target tuning amid distractors. Further, simulations from a restricted model wherein facilitation is purely additive and suppression is purely divisive captures some of the impaired discriminability under crowding in V1, but fails to capture the strength of effects in V4 (Figure 6 – figure supplement 1C-D). Thus, both additive and multiplicative changes together are required to describe how target tuning is altered in V4 and to explain the strong information loss we observe in those population responses.

Beyond these general forms of response modulation under crowding, we showed that V4 tuning for targets is more sensitive to the stimulus configuration than that in V1. V4 neurons have been shown to be selective either for elongated contours (Chen et al. 2014b) or for the curvature of an object within their receptive fields (Pasupathy and Connor, 2002; Nandy et al., 2013; El-Shamayleh and Pasupathy, 2016; Hu et al. 2020). Additionally, V4 neuronal responses depend on the joint stimulus statistics shown within their receptive fields, as in studies of selectivity for stimulus textures (Okazawa et al., 2017). Together, these studies suggest V4 neurons encode the relationship between spatially adjacent image features, in order to extract mid-level representations of object shape and surfaces (Desimone and Schein, 1987; Ghose and Ts’o, 1997; Hegde and Van Essen, 2006; Roe et al., 2012; Pasupathy et al., 2020). While each of these studies probes distinct forms of selectivity, together they reinforce the notion that V4 tuning might differ strongly for a small target stimulus within the receptive field when it appears in isolation or amid other nearby stimuli. Indeed, Motter (2018) showed that V4 neuronal responses to letter stimuli were strongly altered when the arrangement of the constituent line segments was changed. To the extent that downstream areas implement a fixed readout of V4 activity across displays, this context-dependent tuning will limit the resolution of target features available for perception.

Our experiments involved targets and distractors of a fixed size. Would our results have been different if the displays were scaled down so that, for instance, multiple image features fell within individual V1 receptive fields? Crowding effects at a given eccentricity scale with the spacing between target and distractors (Bouma, 1970); when stimuli are small relative to V1 receptive field sizes, discrimination performance can be strongly impaired (Parkes et al., 2001). It remains unclear whether information loss in such displays would be stronger in early cortical areas, or instead remain more prominent in areas like V4. In contrast, if displays were scaled up such that target stimuli were similar in size to the receptive fields of V4 neurons, we would expect the response changes we observed in V1 and V4 to be greatly diminished, consistent with weaker perceptual crowding effects from distractors at increased spatial separation from the target (Bouma, 1970).

Our results convey a more complex picture of how spatial integration operates in cortical populations, one that stands in seeming contrast to simpler descriptions of stimulus features being ‘averaged’ (Parkes et al., 2001) or ‘pooled’ within a single cortical receptive field as an explanation of information loss under crowding. Often these terms are used to indicate a combination or interaction of features, without a precise specification of the mechanistic form of spatial pooling in individual neurons. In this study, we have specified the form that this combination takes, in order to link information loss to the well-studied spatial integration properties of neurons at different levels of the visual hierarchy. It is worth noting that studies that model complex feature selectivity as a joint response dependence among V1-like units, whether physiologically (Freeman et al., 2013) or perceptually (Keshvari and Rosenholtz, 2016), do not incorporate any of the known spatial nonlinearities in V1 responses in their models (e.g., divisive surround suppression). Recent work has shown that the incorporation of these nonlinearities is critical for understanding spatial contextual effects on perception, including crowding (Henry and Kohn, 2020; Ziemba and Simoncelli, 2021).

Previous work has suggested that crowding occurs in V1, V2, or V4, or that it arises from interactions across the visual hierarchy (Freeman et al., 2011b). What do our results imply for where the bottleneck responsible for crowding resides? Using an optimal linear readout of neuronal population activity, we found moderate information loss in V1 and more pronounced loss in V4. We have previously shown that perceptual crowding in our displays, in humans and monkeys, produces a threshold elevation of roughly 50% (Henry and Kohn, 2020), slightly greater than that evident in our V1 data and weaker than that evident in V4. Additionally, perceptual crowding occurs when the spacing between targets and distractors is within roughly half the target eccentricity (Bouma, 1970); we found substantial information loss in V4 for distractors placed farther away than this distance (2.6 deg for target eccentricity of roughly 3 degrees).

Naively, one might be tempted to conclude that crowding arises at some stage between V1 and V4, such as V2. However, discriminability is a function of both population information and the readout strategy. Observers have been shown to employ suboptimal readout strategies (Beck et al., 2012; Jin et al., 2019). Although linear decoders of V1 responses generalize fairly well across the different display configurations we used (Figure 8), deploying slightly suboptimal strategies for reading out V1 responses could easily lead to stronger threshold elevation. Similarly, a suboptimal readout of V4 responses to targets presented in isolation might alter the degree of information loss there. Indeed, many aspects of crowding may be attributable to mechanisms of sensory readout. Attention (He et al., 1996), substitution of distractor features for that of the target (Ester et al., 2015), and perceptual averaging of nearby image features (Parkes et al., 2001) may all reflect sensory readout strategies that are either fixed and suboptimal or fluctuate trial-to-trial.

Together, this suggests a conceptual framework whereby crowding does not reflect a unitary bottleneck on spatial feature perception at a defined locus within the visual system, but instead the result of a series of diverse transformations in visual processing. Visual signals are encoded in cortical populations, then repeatedly re-coded in novel representational formats downstream. Those representations, affected by mechanisms of spatial integration both within each neuron’s receptive field and beyond, are then interpreted through the lens of a readout strategy to determine perception in cluttered, natural environments.

## METHODS

### Subjects and task

We used two male, adult cynomolgus macaques (*Macaca fascicularis*). All procedures were approved by the Institutional Animal Care and Use Committee of the Albert Einstein College of Medicine and were in compliance with the guidelines set forth in the National Institutes of Health *Guide for the Care and Use of Laboratory Animals*.

Animals were first implanted with a titanium headpost to allow for head restraint. Surgery was performed under isoflurane anesthesia, following strict sterile procedures. Antibiotics (ceftiofur or enrofloxacin) and an analgesic (buprenorphine) were provided postoperatively. Animals recovered for a minimum of 6 weeks before behavioral training began.

Animals were trained to enter a primate chair (Crist Instruments) and the head was stabilized via the implanted headpost. Inside a recording booth, animals viewed a CRT monitor (Iiyama; 100 Hz refresh rate; 1024 × 768 screen resolution; 40 cd/m^2^ mean luminance) with linearized output luminance, from a distance of 64 cm. Visual stimuli were generated using custom software based on OpenGL libraries (Expo). Eye position was monitored using a video-based eye-tracking system (SR Research) running at 1 kHz sampling rate.

To initiate a trial, animals had to direct their gaze to a small central fixation point (0.2 degree diameter) and maintain fixation within a 1.3 degree diameter window around that point for the duration of the trial (eye positions were typically < 0.5 degrees from the fixation point). 0.2 s after fixation onset, a visual stimulus was presented in the lower right visual field for 0.25 s. Stimulus offset was followed by 0.35 s of a blank screen (mean grey), and then a second stimulus shown for 0.25 s. If visual fixation was maintained for an additional 0.15 s after the last stimulus offset, animals were given a small liquid reward. Eye movements outside the fixation window led to an aborted trial and no reward; these trials were not analyzed. A typical session consisted of 1000-1300 successful trials.

### Neurophysiological recording

Once fixation training yielded stable behavioral performance, we performed a second sterile surgery, making a craniotomy and durotomy in the left hemisphere of each animal and implanting microelectrode arrays (Blackrock Microsystems) into V1 and V4. Array placement was targeted to yield overlapping retinotopic locations in the two areas, based on anatomical landmarks and prior physiological studies (Van Essen et al., 1984; Gattass et al., 1988). Each electrode array had either a 10 × 10 or a 6 × 8 configuration. In one animal, we implanted 1 array in V4 (48 channel) and 2 in V1 (48 and 96 ch). In the other, we implanted 2 arrays in V4 (both 48 ch) and 2 in V1 (both 96 ch). After insertion, we sutured the dura over the arrays and covered it with a matrix to promote dural repair (DuraGen; Integra LifeSciences). The craniotomy was covered with titanium mesh.

Extracellular voltage signals from the arrays were amplified and recorded using either Cerebus (Blackrock Microsystems) or Grapevine Scout (Ripple Neuro) data acquisition systems. Signals were bandpass filtered between 250 Hz and 7.5 kHz, and those exceeding a user-defined threshold were digitized at 30 kHz and sorted offline (Plexon Offline Sorter). The local field potential on each electrode was also recorded but not analyzed in this study.

### Visual stimuli

In initial sessions, we mapped the neuronal receptive fields. We presented small grating stimuli on a 9 × 9 degree grid of locations (sampled at 1 degree resolution; each stimulus shown at 4 orientations separated by 45 degrees, luminance contrast 50 or100%, stimulus size 0.5-1.0 degrees). Using the resultant maps, we then placed the target stimuli so as to drive both the V1 and V4 populations. We also conducted a brief, coarse mapping of each neuron’s receptive field at the beginning of each recording session.

Target stimuli were drifting sinusoidal gratings (spatial frequency of 1.5 cycles per degree, temporal frequency of 4 Hz, 1 degree diameter) of 50% luminance contrast. For each session, a reference orientation was chosen, and target orientations were sampled from an 18-degree range spanning that reference. On a subset of trials, 4 distractor stimuli (of the same size and spatiotemporal frequency as the target) were shown adjacent to the target stimulus. The orientation of the distractors was fixed at +/-10 degrees from the reference, and tilt was spatially alternated clockwise around the target. All stimuli had simultaneous onset and offset, and started their drift from a fixed phase. Across trials, the center-to-center spacing of distractors from the target was varied from 1.3 to 2.6 degrees. A second set of stimuli, identical to the first except for a 90-degree rotation of the ensemble reference orientation, was included in each session, to mitigate adaptation effects from repeated presentation of one range of orientations. All stimuli (targets alone, targets with distractors, and changes in distractor spacing and reference orientation) were randomly interleaved. In a separate set of experiments (N = 4 sessions, reported only in Figure 7E), we fixed the spacing of the distractor stimuli but instead varied their orientation, drawing from two distractor arrays with counterbalanced orientations.

### Data analysis

Analysis was restricted to neurons that were driven by the target stimuli in isolation (responses ≥ 1 s.d. above spontaneous rate). All analyses were performed on the total spike count of each neuron during stimulus presentation on each trial (from 0-0.25 s after stimulus onset).

Discriminability of target orientation in single neuron responses was quantified by applying receiver operating characteristic analysis (Green and Swets, 1966) to the spike count distributions evoked from presentations of the two most extreme target orientations shown (18 degree separation). We used the area under the ROC curve as the measure of discriminability. We used the slope of the regression line relating discriminability for targets alone to discriminability in the presence of distractors to quantify the average change in discriminability across neurons. 95% confidence intervals for the slope were estimated using bootstrap with replacement across neurons.

Target orientation discriminability in neuronal populations was quantified using logistic regression, with Lasso regularization. The regularization parameter was set to the largest value that yielded maximal cross-validated performance on held out trials from the training set. For each pairwise combination of target orientations shown (and each distractor spacing), we fit a decoding model and evaluated performance on held out test data. The reported decoding performance is an average over cross-validation folds.

Crowding effect size for a given target-distractor separation was estimated by combining decoding performance values for target alone and crowded conditions across pairings of target stimuli and across recording sessions. We fit a single parameter model to the combined data to characterize the threshold elevation under crowding that best explained the change from target alone to crowded performance, using maximum likelihood estimation (validation on synthetic data is shown in Figure 3 – figure supplement 1). Specifically, a target only discrimination model consisted of a cumulative normal distribution adjusted for chance performance in the 2AFC task. For each population and target only performance, we found the stimulus that yielded this performance by inverting the discrimination model. Then, for a given model threshold elevation under crowding, we could generate a predicted discrimination performance. This predicted model performance, assuming a binomial distribution, yielded a probability of the observed discrimination performance under crowding given this model of threshold elevation. For each model, we combined data points by taking the product of likelihoods, and threshold elevation was estimated to be the model with the maximum likelihood. We used a nonparametric bootstrap procedure (sampling from the data points with replacement) to generate confidence intervals for these estimates.

Spike count correlations (r_SC_) were computed using trials in which the response of either neuron was within 3 s.d. of its mean response, to avoid contamination by outlier responses (Kohn and Smith, 2005). Correlation values were calculated separately for each target orientation and then averaged.

We assessed how the target orientation tuning of each neuron changed with distractors by finding the parameters *m* and *a*, that minimized the difference between the measured responses to targets amid distractors and the predicted responses. The predicted responses, *r*_*TD*_, were given by 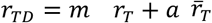, where *r*_*T*_ are the given responses to isolated targets, and 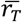 is the mean response to isolated targets. We performed this analysis on neurons whose tuning was consistent, defined by calculating the correlation between the target-only tuning computed on even versus odd trials. Neurons with a correlation above 0.5 were included.

To assess the similarity of the population representation for targets under different crowding conditions, we compared population decoding performance across conditions using a shared, fixed decoding strategy. We fit linear decoders with regularization, as previously described, to the population responses to the two most differing target orientations. One decoder was estimated using responses to targets in isolation, and a second decoder (‘crowded’) was estimated from responses to target stimuli with nearby distractors. Responses on held out trials were projected onto these two readout axes, as were an equal number of trials from the other target orientations and conditions of crowding. Population discriminability for two target stimuli along a specific readout axis was quantified by the d-prime between the response distributions along the decoding axis, where 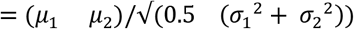. We compared d-prime measures for responses projected onto the readout axis for the matching condition (‘same decoder’) to those projected onto an opposing crowding condition (‘cross decoder’).

All indications of variance in the text and figures are SEM unless otherwise noted.

## Acknowledgements

We thank Amir Aschner and Anna Jasper for assistance with experiments. This work was supported by NIH grants (EY023926 and EY028626) and the Charles H. Revson Senior Fellowship in Biomedical Science.

## Author Contributions

CAH and AK designed the project; CAH performed experiments and analyzed the data; CAH and AK wrote the paper.

---

**Figure 3 – figure supplement 1.**
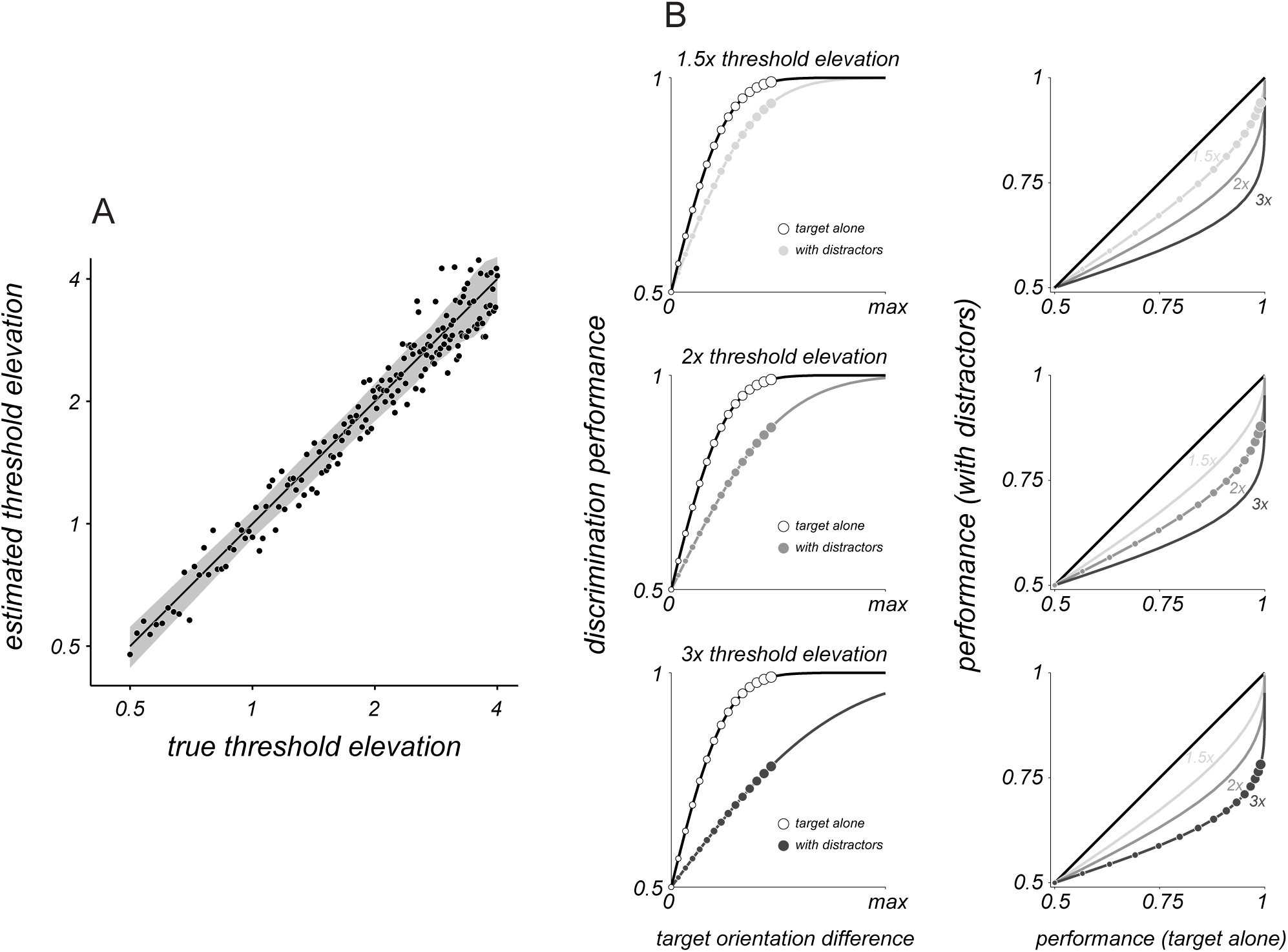
Method to estimate threshold elevation. A. Ground truth change in discrimination under crowding was modeled for a range of elevations of discrimination threshold compared to targets alone (abscissa). Empirical threshold elevation was inferred from the simulated data (n = 40 trials per stimulus) by finding the single parameter change in threshold via maximum likelihood given the data. Black line indicates the unity line, shaded gray region represents ±1 s.e.m. B. Plots of discrimination performance for distinct threshold elevations under crowding. Left column: Performance for targets alone (white) and with distractors (gray) as a function of orientation difference. Right column: Target alone performance (abscissa) is plotted in relation to performance with distractors (ordinate). Gray curves indicate expected relationship for crowding models with 3 different changes in threshold. Rows indicate models with progressively greater threshold elevations under crowding.

**Figure 3 – figure supplement 2.**
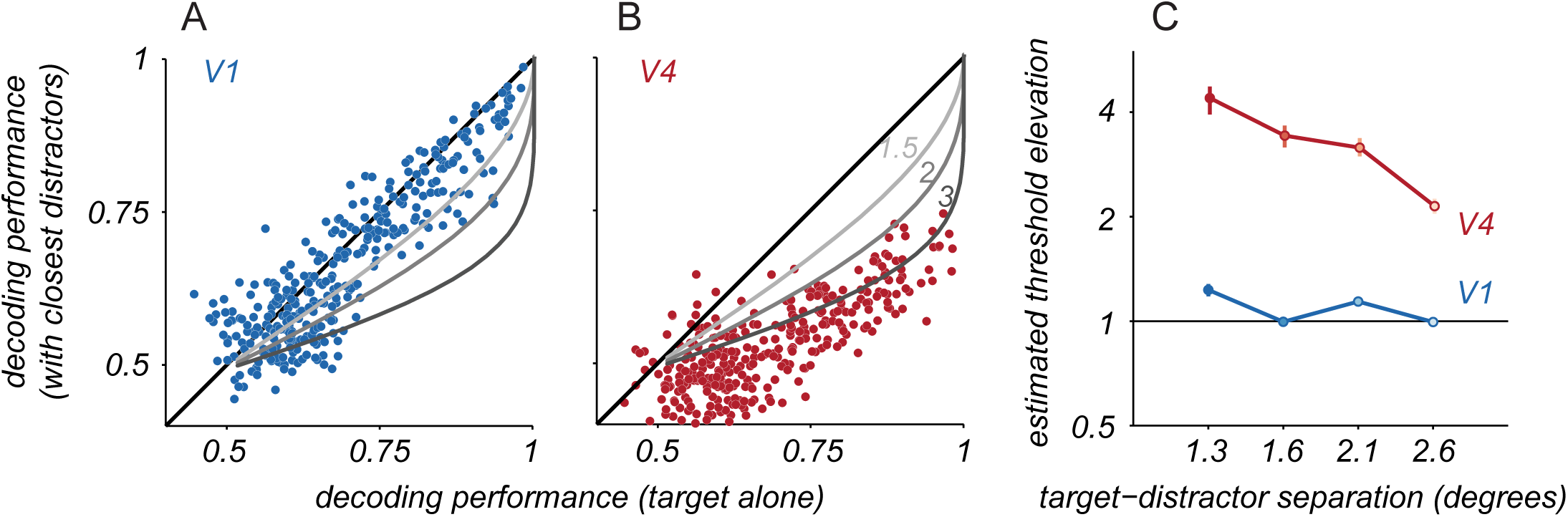
Target discriminability in pseudo-populations. A. Pseudo-population responses were created for a range of population sizes by resampling from neuronal responses (across animals and sessions). Cross-validated V1 pseudo-population decoding performance for targets alone (abscissa) and targets with closest distractors (ordinate). Each point represents a single pairing of two target stimuli. Gray lines indicate expected performance for the indicated threshold elevations. B. Same as in A, but for V4 pseudo-populations. C. Threshold elevation as a function of target-distractor separation. Points and error bars represent the mean ± s.e.m. estimated via bootstrapped resampling of the data for V1 (blue) and V4 (red) pseudo-populations.

**Figure 6 – figure supplement 1.**
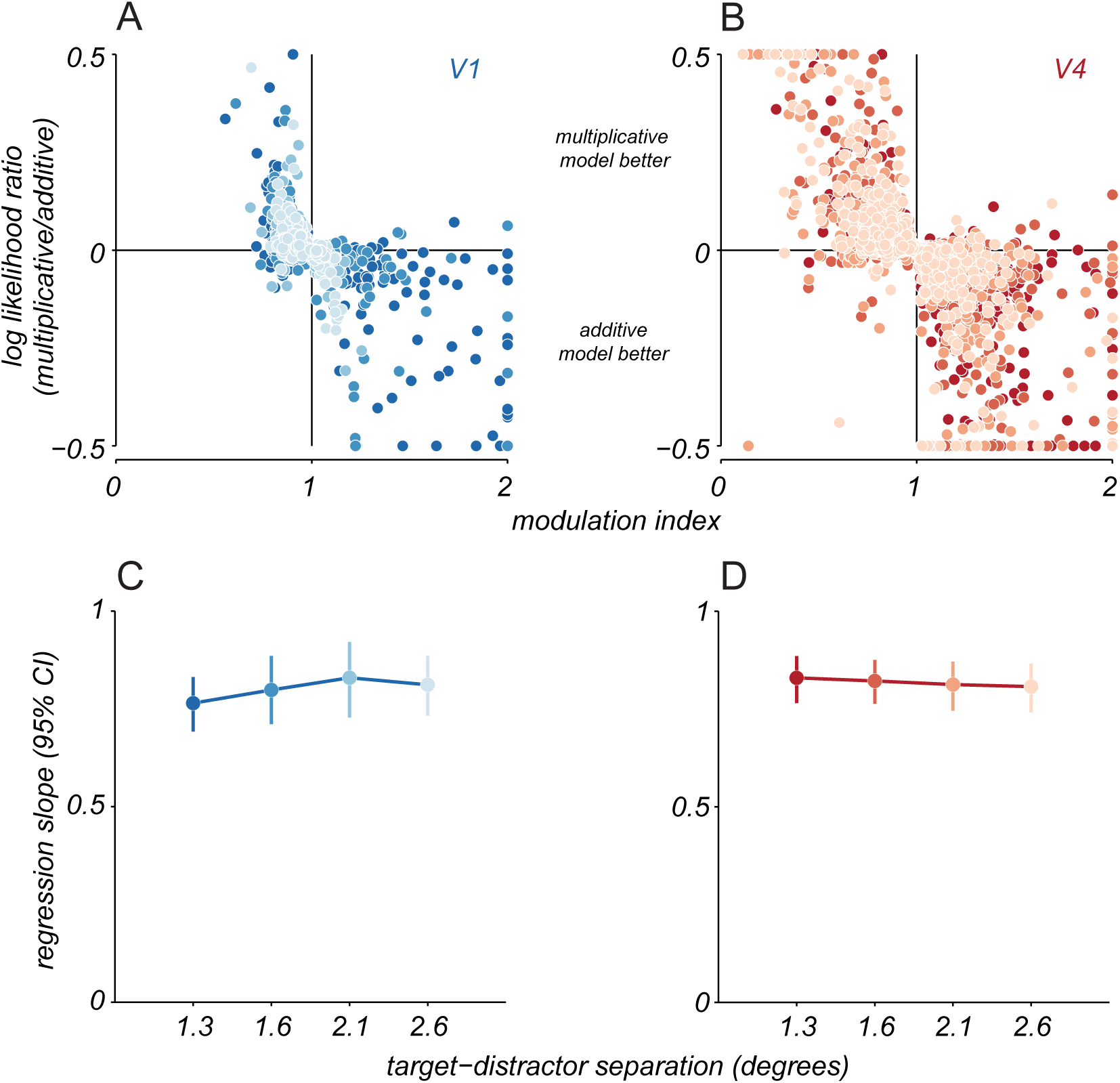
Comparison of additive and multiplicative modulation of responses to targets by distractors. We repeated an analysis from our prior study (Henry and Kohn, 2020) to determine whether the orientation tuning for targets amid distractors was better characterized as an additive or multiplicative change of the tuning for isolated targets. Responses under crowding were fit by inclusion of either an additive or multiplicative scaling term c, applied to the responses to targets alone. Model responses were rectified at 0. Model comparison was performed using the log-likelihood of each model given the data, assuming Poisson variability of neuronal responses (El-Shamayleh and Movshon, 2011). Note that our definition of additive modulation of tuning does not require that responses to targets and distractors strictly add. Sublinear (e.g. due to normalization, Carandini and Heeger, 2011) or supralinear (e.g. due to thresholding or exponentiation, Carandini et al., 2005) summation would also result in an additive modulation of tuning, so long as the degree of non-linear summation was constant across target orientations. A.Log-likelihood model comparison for whether V1 tuning with distractors was better explained by an additive or multiplicative change in tuning for targets alone. Response modulation index is shown on the abscissa; values less than 1 indicate suppression, greater than 1 indicate facilitation. Each point represents a single V1 neuron and distractor spacing condition (near distractors: dark blue, far: light blue). B. Model comparison for V4 neurons, same conventions as in A. In both areas, an increase in responsivity (modulation index>1) was better described as an additive modulation; a loss of responsivity (modulation index<1) was better described as a multiplicative modulation. C. Both additive facilitation and multiplicative suppression would be expected to reduce discriminability (Henry and Kohn, 2020). To determine how strongly discriminability would be affected, we simulated responses for each neuron using a model where facilitation acts additively and suppression divisively. This panel shows the expected change in V1 discriminability as a function of target-distractor spacing. The change in discriminability was quantified using the slope of the regression line relating discriminability for target only and crowded conditions, as in Figures 2 and 6. Error bars indicate 95% confidence interval. D. Same as in C, but for V4 discriminability. Thus, panels C and D show that allowing for either additive facilitation or divisive suppression reduces discriminability in both areas, but not as strongly as in the measured responses. Only by including both forms of modulation together (Figure 6) can we accurately capture the observed change in discriminability.

**Figure 8 – figure supplement 1.**
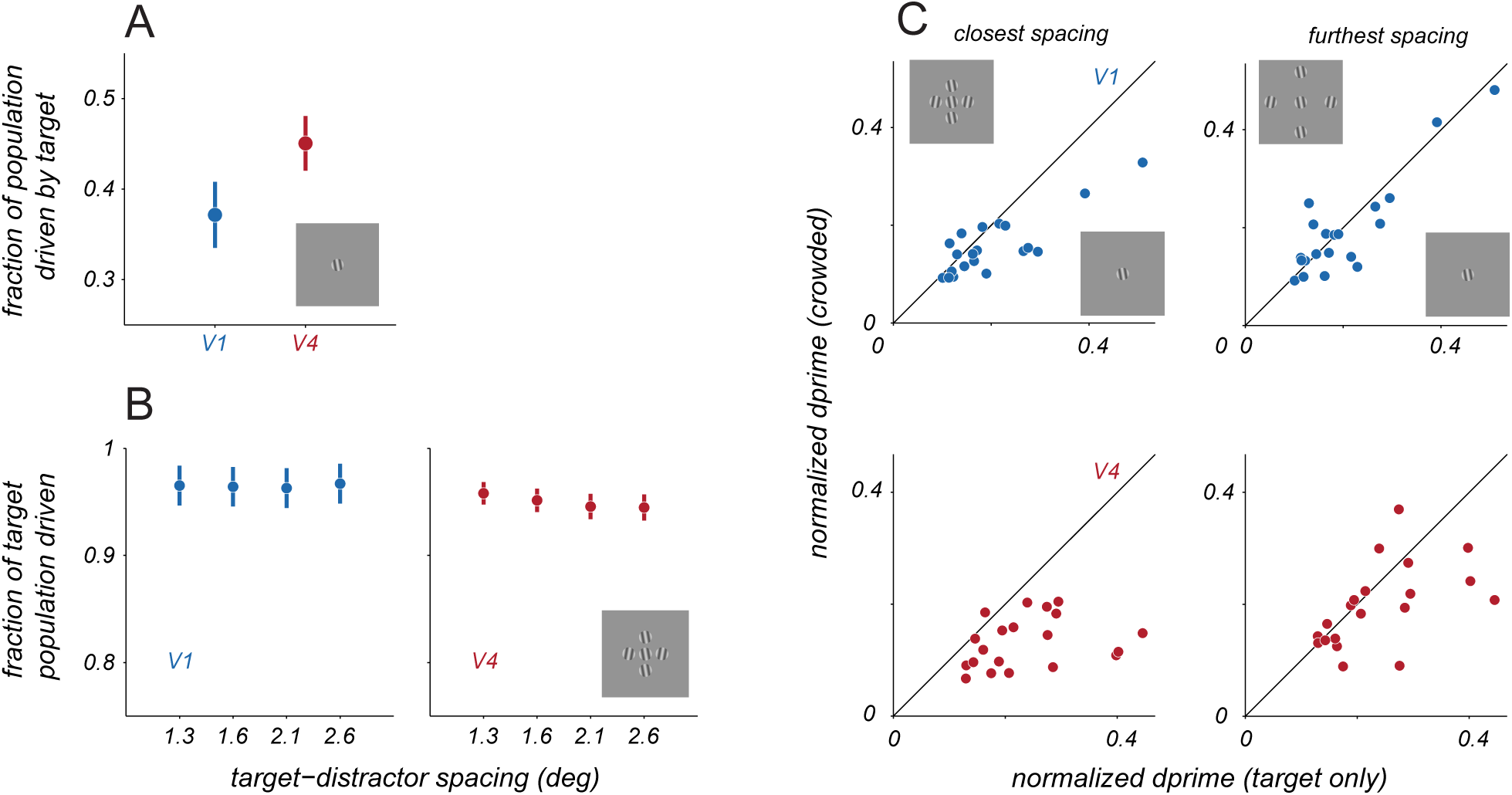
Informative populations across conditions of crowding. A. Average fraction of neuronal population that was significantly driven by target stimuli (defined as a response greater than the baseline rate and 1 SD of that rate), for V1 (blue) and V4 (red). Points represent mean ± s.e.m. B. Average fraction of those target-responsive populations that remained driven in the presence of distractors, at various target-distractor spacings. C. For each population, the contribution of the most informative neuron for targets alone (abscissa) compared to the informativeness of that neuron for targets with distractors (ordinate). Dprime values were calculated for the two most extreme targets shown, and then normalized by the summed dprime value of the recorded population. Left column represents comparison between targets alone and with near distractors; right column represents comparison between targets alone and with far distractors.

